# Structure of the Bacteriophage PhiKZ non-Virion RNA Polymerase

**DOI:** 10.1101/2021.04.06.438582

**Authors:** Natàlia de Martín Garrido, Mariia Orekhova, Yuen Ting Emilie Lai Wan Loong, Anna Litvinova, Kailash Ramlaul, Tatyana Artamonova, Alexei S. Melnikov, Pavel Serdobintsev, Christopher H. S. Aylett, Maria Yakunina

## Abstract

Bacteriophage ΦKZ is the founding member of a family of massive bacterial viruses. It is considered to have therapeutic potential as its host, *Pseudomonas aeruginosa*, is an opportunistic, intrinsically antibiotic resistant, pathogen that kills tens of thousands worldwide each year. ΦKZ is an incredibly interesting virus, expressing many systems the host already possesses. On infection, it forms a “nucleus”, erecting a barrier around its massive genome to exclude host restriction endonucleases and CRISPR-Cas systems. ΦKZ infection is independent of the host transcriptional apparatus. It expresses two different multi-subunit RNA polymerases (RNAPs): the virion RNAP (vRNAP) is injected with the viral DNA during infection to transcribe early genes, including those encoding the non-virion RNAP (nvRNAP), which transcribes all further genes. ΦKZ nvRNAP is formed by four polypeptides thought to represent homologues of the eubacterial β/β′ subunits, and a fifth with unclear homology, but essential for transcription. We have resolved the structure of ΦKZ nvRNAP to 3.3 Å, shedding light on its assembly, homology, and the biological role of the fifth subunit: it is an embedded, integral member of the complex, with structural homology and a biochemical role implying that it has evolved from an ancestral homologue to σ-factor.

## Introduction

Bacteriophage ΦKZ is the prototypical member of a family of giant bacterial viruses distantly related to the *Myoviridae* (1, 2). It infects the intrinsically antibiotic resistant opportunistic pathogen *Pseudomonas aeruginosa*, which kills tens of thousands annually, and is therefore considered a potential candidate for medical use as a bacteriophage therapy (3, 4). ΦKZ is of exceptional size with a massive DNA genome (280 334 bp)(5) encoding many proteins that the host would typically be expected to provide during infection, and several other systems that the host lacks entirely. During infection, ΦKZ forms a “nucleus” within the host cytoplasm that protects its genome from host endonucleases, and which it traffics with a tubulin cytoskeleton (6, 7), providing intriguing evolutionary parallels to the eukaryotic nuclear machinery. In many ways, bacteriophage ΦKZ has been found to be more complex and scientifically interesting than the organism it infects.

The lifecycle of ΦKZ in *P. aeruginosa* is unaffected by the host RNA polymerase (RNAP) inhibitor rifampicin, indicating that ΦKZ is independent of the host transcription machinery as it provides its own RNA polymerases (8). Whereas almost all known bacteriophage-encoded RNA polymerases are single subunits, ΦKZ is one of only a few viruses (9–11) to encode multisubunit RNAPs (msRNAPs). All known msRNAPs comprise the conserved double-ψ β-barrel (DPBB) domain in the two largest subunits, β and β’ in bacteria (12). These largest subunits are usually catalytically inactive without auxiliary subunits. In eubacteria, a dimer of α subunits and a single ω subunit are required to assemble the minimal catalytically active core capable of the RNA polymerisation reaction (ββ’α_2_ω)(13–16). Although this core is catalytically active, exchangeable σ-subunits are required to initiate transcription with specificity from their cognate promoters at the correct transcription initiation sites (17, 18).

Bacteriophage ΦKZ encodes two sets of proteins homologous to small subsections of the two largest subunits of bacterial RNAP, β and β’, but no known homologues of the α or ω subunits usually required for transcription at all; a ΦKZ virion RNAP is formed from at least four polypeptides: gp178, gp149, gp180, and gp80, and is injected with the genome to express ΦKZ early genes (8), while a ΦKZ non-virion RNAP (nvRNAP) is formed from these early genes by five polypeptides: gp55, gp68, gp71-73, gp74, and gp123 (10). Both gp55 and gp74 exhibit a degree of homology to the β’ subunit, whereas gp71-73 and gp123 represent similarly weakly homologous counterparts of the β subunit. No homology had previously been identified for ΦKZ-gp68 (10), however *in vitro* transcription assays have revealed that complexes lacking gp68 are unable to elongate existing RNA or bind to a DNA-RNA scaffold, demonstrating that gp68 is essential for even the most basic biological functions of the complex (19).

In this study, we have determined the three-dimensional structure of ΦKZ nvRNAP to 3.3 Å using cryogenic electron microscopy (cryo-EM), allowing atomic modelling of the majority of the complex. Our structure confirmed the previously predicted homology of the split β/β’-like subunits, but further revealed that subunit gp68 is integral to the ΦKZ nvRNAP complex, interacting with both gp71-73 and gp55, and biochemical studies of nvRNAP assembly implied that Gp68 plays a decisive role in the formation of the active centre. We used DNA binding assays to demonstrate that gp68 is essential for both promoter binding and recognition, and structural comparison with other RNAPs showed that it has homology to, and is located in the same place as, bacterial σ factors.

## Materials and Methods

### Cloning

The set of pET-based plasmids containing one of the phiKZ nvRNAP subunits genes, the pnvCo-Ex plasmid, and the pGp68 plasmid, were generated previously (19). PACYC184 was digested using SalI, and blunt ends generated using Klenow fragment (Thermo Fisher Scientific, Waltham, MA, USA). The set of pACYC-based plasmids was created by blunt end insertion of PCR-products with sequences corresponding to the T7 promoter, lac operator, ribosome binding site, and requisite genes of the phiKZ nvRNAP subunit without an additional His-tag from the pET-set of plasmids into pACYC184. pET19K-Hisgp55-(T7pr_gp71-73), or pET28a-Hisgp68-(T7pr_gp71-73) plasmids were assembled from two PCR-fragments using the NEBuilder HiFi DNA Assembly Master Mix (New England Biolabs, Ipswich MA, USA) according to the manufacturer’s protocol.

### Protein expression and purification

BL21(DE3) *E. coli* cells were transformed with either one plasmid, or two plasmids with compatible origins. Expression was induced by the addition of 1 mM IPTG once an OD_600_ of 0.5-0.7 had been reached, and cultures were incubated at 22 °C for 3 h after induction. Recombinant ΦKZ nvRNAP complexes were purified as previously described (19).

For subunit complex validation, 1 g wet mass of each corresponding cell pellet was disrupted by sonication in 10 mL of buffer A (40 mM Tris-Cl pH 8.0, 10% glycerol, 500 mM NaCl,1 mM DTT) containing 5 mM Imidazole followed by centrifugation at 11000 rcf for 30 min. Clarified lysate was loaded onto a HisTrap HP 1 mL (GE LS, Chicago IL, USA) that had previously been equilibrated, and then washed with buffer A with 5 mM Imidazole. The recombinant complexes were then eluted with buffer A containing 250 mM Imidazole. Size-exclusion chromatography of the recovered factions was performed over a Superdex 200 Increase 10/300 GL (GE LS, Chicago IL, USA) in TGED buffer (20 mM Tris-Cl pH 8.0, 5% glycerol, 0.5 mM EDTA, 1 mM DTT) with 200 mM NaCl.

For purification of Gp68 without a tag, 1 g wet mass of cell pellet was resuspended in 10 mL TGED buffer with 50 mM NaCl and disrupted by sonication. Clarified lysate was passed through a HiTrap Q XL column (GE LS, Chicago IL, USA) equilibrated into the same buffer. The resulting sample was loaded onto HiTrap Heparin HP (GE LS, Chicago IL, USA) followed by washing using TGED buffer with 100 mM NaCl. The protein was eluted using TGED buffer with 200 mM NaCl. The eluted fractions were concentrated (Amicon Ultra-4 Centrifugal Filter Unit with Ultracel-30 membrane, EMD Millipore, Merck, Burlington MA, USA) and purified by size exclusion chromatography using a Superdex 200 Increase 10/300 (GE LS, Chicago IL, USA) in TGED buffer with 200 mM NaCl.

### DNA-template preparation

The RNA-DNA scaffold and all types of DNA templates were prepared as previously described (Orekhova et al., 2019). The oligonucleotide primers used are listed in Supplementary Table 4.

### Native PAGE and in vitro transcription

Native polyacrylamide gel electrophoresis and *in vitro* transcription reactions were performed as previously described (19).

### Electrophoretic mobility shift assay (EMSA) and photo-crosslinking

Electrophoretic mobility shift assay (EMSA) was carried out as previously described (19). For photo-crosslinking reactions the reaction mixture (10 μL) containing 5 pmol of the template DNA and 7.5 pmol of ΦKZ nvRNAP were incubated in 1x transcription buffer (40 mM Tris-Cl pH 8.0, 10 mM MgCl_2_, 5 mM DTT) for 15 minutes at 37 °C. To induce the formation of covalent cross-links, the mixture was irradiated for 15 seconds. A RAPOP-100 femtosecond laser system and an ATsG800-17 third harmonic generator (“Avesta-Project”) were used as a source of ultraviolet radiation with a wavelength of 266 nm. The radiation parameters (the 3rd harmonic) were; radiation pulse energy of 10 μJ, pulse duration of 80 fs, repetition frequency of 2 kHz, and beam diameter of 2 mm. The resulting average power was 20 mW, and the radiation power density in the pulse was 4 GWcm^-2^. To break down unsuccessfully crosslinked complexes, heparin was added to the reaction mixture to a concentration of 100 g/L and reaction mixture was incubated at 37 °C for 10 minutes. Samples were analyzed by EMSA in DNA loading buffer (8.3% glycerol) or SDS-DNA loading buffer (8.3% glycerol, 0.2% SDS, 30 mM EDTA) before loading onto the 8% TBE-polyacrylamide gel.

### Limited proteolysis

ΦKZ nvRNAP (10 pmol), and its complexes with DNA-templates (20 pmol), were digested using trypsin (Sigma-Aldrich, St. Louis, MO, USA) in 100 mM Tris-Cl pH 8.5 at 22 °C for 30 min. The concentration of trypsin used was 0.09 g/L. The proteolysis reactions were quenched through the addition of SDS-PAGE sample buffer, followed by immediate incubation of each sample at 100 °C for 5 minutes. The resulting samples were analyzed by SDS-PAGE (8 and 10% polyacrylamide gels).

### Mass-spectrometry

Protein bands of interest were manually excised from the Coomassie-stained SDS or native TBE-PAGE. Individual slices from SDS or native PAGE were prepared for mass-spectrometry through *in situ* trypsin digestion at 37 °C for 4h as previously described (10). In the case of slices of TBE-PAGE gels, no washing step was performed.

### Grid preparation

ΦKZ nvRNAP complexes were exchanged into sample buffer containing 15 mM Tris-Cl pH 8.0, 150 mM NaCl, 0.5 mM EDTA, 2 mM MgCl_2_, 1 mM DTT, to a final concentration of 0.3 mg/mL. Samples of ΦKZ nvRNAP were adsorbed to a thin film of graphene oxide deposited upon the surface of holey carbon copper grids (R2/1, 300 mesh, Quantifoil). Grids were blotted for 1-2 seconds before plunge freezing in liquid ethane using a Vitrobot Mark IV (Thermo Fisher Scientific, Waltham, MA, USA) at 4 °C and 100% humidity.

### Calculation of reference density

A total of 257 micrographs from a cryo-grid of ΦKZ nvRNAP were collected on an early Falcon direct electron detector using an FEI Tecnai F20 electron microscope (Thermo Fisher Scientific, Waltham, MA, USA) at a magnification of 81000-fold, an acceleration voltage of 200 kV, and a total dose of 50 e^-^/Å^2^ over a defocus range of -2 to -5 μm. A dataset of 45 980 particles was selected semi-automatically using BOXER (28). The parameters of the contrast transfer function were determined using CTFFIND4 (29). Particles were 2D-classified into 100 classes in two dimensions using RELION 3.0 (30) and 7 well-defined classes including 18 508 particles were selected for initial three-dimensional reconstruction. Initial models were generated using the stochastic gradient descent approach in RELION 3.0, then filtered to 50 Å and used as an initial reference for automatic refinement. Projections from the resulting initial models were consistent with class-averages, and were therefore used for further refinement of higher resolution data.

### Dataset acquisition

Data from ΦKZ nvRNAP samples was collected on a Titan Krios G3i (Thermo Fisher Scientific, Waltham, MA, USA) at the London Consortium for Electron Microscopy microscope sited at the Crick institute, equipped with a K3 direct electron detector (GATAN, San Diego, USA) and operated at 300 kV, 37 000-fold magnification and with an applied defocus range of -0.75 to -3.25 μm. Frames were recorded automatically using EPU, resulting in 10 582 images of with a pixel size of 1.1 Å on the object scale. Images were recorded as stacks of 30 separate frames in electron counting mode, comprising a total exposure of ∼40 e^-^Å^-2^.

### Data processing

Frames were aligned, summed and weighted by dose according to the method of Grant and Grigorieff using MotionCor2 to obtain a final image. Poor-quality micrographs were rejected based on diminished estimated maximum resolution on CTF estimation using CTFFIND4 and visually based on irregularity of the observed Thon rings. Particles were selected using BATCHBOXER (31), and refinement thereafter performed using RELION 3.0.

Two-dimensional reference-free alignment was performed on 3 574 387 initial particles to exclude those that did not yield high-resolution class averages. Of these, 1 945 924 particles were retained for further refinement. These particles were initially refined to high resolution using the auto-refinement procedure, in RELION 3.0, then classified into 6 classes in 3-dimensions, the highest resolution class with the largest ordered fraction of the molecule being retained for final high-resolution refinement. This final gold-standard refinement of ΦKZ nvRNAP from 427 214 particles reached 3.3 Å according to an independent half-set FSC of 0.143.

### Modelling and refinement

Initial models for the most homologous regions of the ΦKZ nvRNAP were prepared using Swiss-Model based on PDB-6EDT, and FUGUE. The model was then rebuilt with COOT and extended outwards from these well-defined reference points to the remaining regions of the four homologous proteins. ΦKZ-GP68 was finally built into the remaining density, initially as a poly-alanine model, before the sequence register was eventually established based upon the identification of clear patterns of large side-chains and secondary structure elements (32). The atomic model was refined with PHENIX real-space refine. Homology searches and Cα comparisons were carried out using the DALI-lite server, while surface area calculations were performed using the PISA server.

## Results and Discussion

We generated recombinant ΦKZ nvRNAP complexes in *Escherichia coli* using a co-expression system with one plasmid bearing the four β/β’-like proteins and another encoding gp68 to allow simple generation of both four and five-polypeptide ΦKZ nvRNAP, as well as free gp68, for biochemical experiments (Supplementary Figure 1). For structure determination, ΦKZ nvRNAP complexes were stabilised by adsorption to a graphene oxide film covering a holey carbon copper grid and vitrified before visualisation by cryo-EM. Electron micrographs of vitrified ΦKZ nvRNA particles proved similar in appearance to eubacterial msRNAP (20), and we resolved the structure by single particle analysis to a final resolution of 3.3 Å (FSC=0.143) (Supplementary Figure 2). Initial comparison of our structure with several previously resolved msRNAPs allowed us to establish positions for the most conserved core regions of the β/β’-like subunits, and we expanded upon these conserved segments, then interpreted the remainder of the density to provide an atomic model covering the majority of the complex (Figure 1, Supplementary table 1).

**Figure 1:**
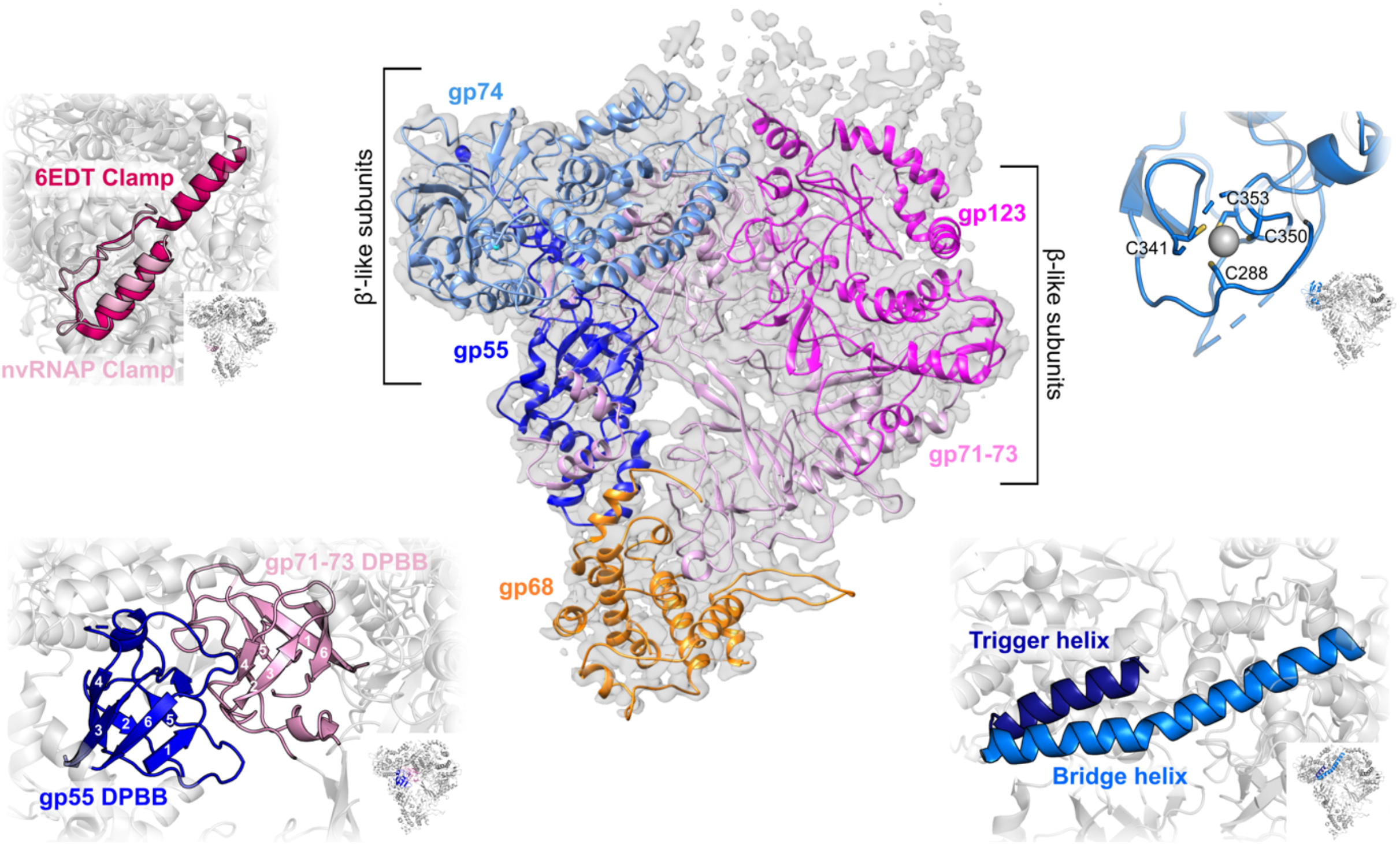
The overall architecture of the ΦKZ nvRNAP is homologous to eubacterial msRNAP. Electron scattering density of the ΦKZ nvRNAP with our molecular model shown in cartoon representation. Insets: (top-left) Superimposition of nvRNAP clamp (light pink) on the *M. tuberculosis* RNAP clamp (dark pink). (top-right) Zn-finger of gp74 (blue). (bottom-left) DPBBs of gp55 (blue) and gp71-73 (pink). Each strand is numbered from N-terminus to C-terminus. (bottom-right) Trigger helix (deep blue) and bridge helix (light blue) of gp74.

The overall architecture of ΦKZ nvRNAP proved to resemble that found in eubacterial msRNAPs, the closest structural homologue being *Mycobacterium tuberculosis* RNAP (PDB: 6EDT)(21) Cα-RMSD 3.2 Å. The ΦKZ nvRNAP complex retains the conserved β/β’ architecture characteristic of msRNAPs, however the overall appearance is more akin to a “clam-shell” than the canonical “crab-claw”, with a shallow active site channel running along the elongated side of the structure (Figure 1). One side of the enzyme is formed by the β’-like subunits, gp55 and gp74, with the β-like subunits gp71-73 and gp123 forming the opposing side, the internal DNA channel running between them through the centre of the complex as expected. Density corresponding to the most peripheral parts of the enzyme which would encircle the channel is weakly resolved in our structure due to flexibility, presumably given the absence of a substrate. Through comparison with the eubacterial RNAP structure, we suggest that the principle flexible region we see belongs to the N-terminus of gp55, which is missing in our model, however sections of gp68, gp74, and gp123 are also too flexible to model (Supplementary Figure 3).

The key active elements and enzymatic residues of ΦKZ nvRNAP are well-conserved, implying an identical reaction cycle to eubacteria, despite the split and missing subunits. The DPBBs that form the catalytic core are contributed by gp55, which contains the universally conserved DxDGD motif (14, 22) that coordinates the catalytic Mg^2+^ (^417^DFDGD^421^ – flexible in our structure) between β5 (403I-M405) and β6 (Q422-L427), and by gp71-73, which contains the two conserved lysines (K386 and K394) believed to be responsible for the interaction with DNA (12)(Figure 1). The structural elements that govern the msRNAP reaction cycle are also conserved in ΦKZ nvRNAP; β’-like subunit gp74 contains the “bridge” helix (L242-Y279) and the “trigger” helix/loop (Figure 1), which switch between alternate conformations during nucleotide addition (22, 23), although only the first helix of the “trigger” loop (residues T365-V383) is well-ordered in our structure, as well as the conserved Zn^2+^ binding domain (C288, C341, C350, C353), known to interact with the promoter during transcription initiation (Figure 1E)(22). The “clamp”, a flexible element whose position is key to regulate the entry of single-stranded DNA into the active site (24), is also conserved and the. visible elements are positioned similarly to their position within the *M. tuberculosis* open promoter complex (RPo) structure (Figure 1)(10), although the remainder of the clamp is only partially ordered at the front of the enzyme within the gp55 N-terminal domain.

The five-subunit ΦKZ nvRNAP complex is sufficient to initiate transcription without α or ω accessory subunits (19). When we assembled full nvRNAP *in vitro* through the addition of independently purified gp68 to 4-subunit (4s) complexes, including only the four β/β’-like subunits, the resulting complexes contained all five nvRNAP components and successfully bound 212 bp specific DNA-templates in a similar manner to *in vivo* 5s assembled complexes (Figure 2A), however, they proved unable to initiate transcription from an RNA-DNA scaffold (Figure 2B). This implied that gp68 is included in the complex at the earliest stage of assembly. Subunit co-purification experiments in which we co-expressed pairs of subunits with one affinity-tag and assayed for recovery of both proteins showed that gp68 bound gp71-73 and gp55 (Supplementary Table 2), which was consistent with our structure in which the fifth subunit, gp68, is integrated into the nvRNAP making extensive contacts with gp71-73, and also contacting gp55 (Figure 1, Supplementary Table 3). The interaction between gp68 and both subunits is mostly governed by hydrogen bonds, with 4% and 14% of the solvent-accessible area of gp68 contributing to interactions with gp55 and gp71-73 respectively. In eubacterial msRNAPs, the dimerised α subunits serve as an organising platform for the assembly of a catalytically active core. Both β and β’ interact with the α-dimer, with universally conserved regions βa14 and βa15 being essential (25). In our structure, βa15 interacts with gp68 (Figure 3A), suggesting that gp68 at least contributes to the stabilisation of the catalytic core, an observation that is consistent with the requirement for gp68 during the assembly process in order to form a catalytically active polymerase.

**Figure 2:**
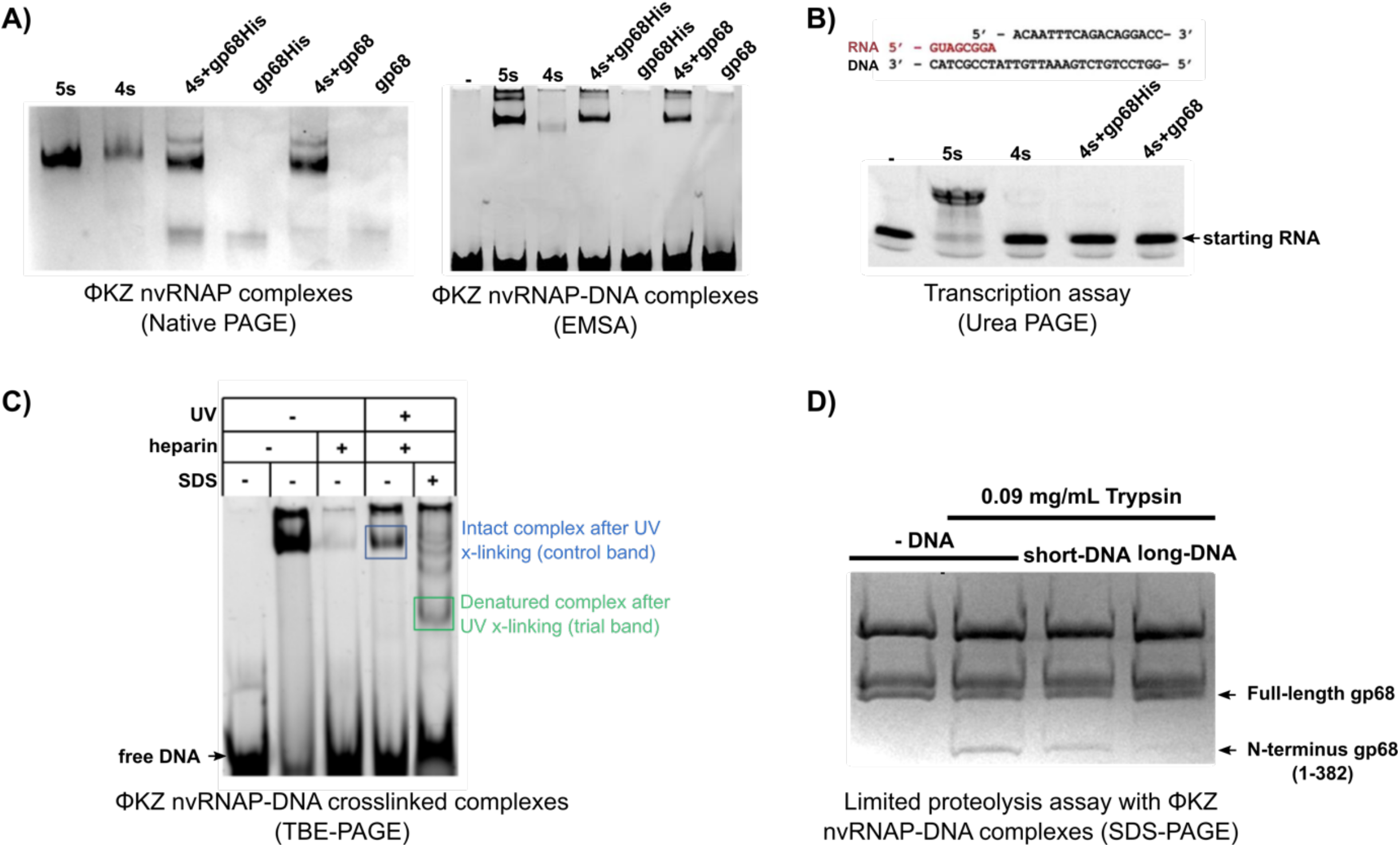
Gp68 is necessary for assembly of active polymerase, and binding of cognate DNA. a) Native PAGE of 4, and 5 subunit complexes prepared through the inclusion of gp68 at different points, and EMSA with a 212 bp specific dsDNA-template containing the KZP119L promoter sequence for different constructs and assemblies of recombinant nvRNAP complexes and independently purified gp68-His and gp68 samples. b) PAGE of the transcription assay from the RNA-DNA scaffold shown above. Activity is indicated by decreased mobility (lane 5s). c) SDS-PAGE of UV cross-linked DNA bound complex, showing bands extracted for subsequent analysis by mass-spectrometry. d) Protection of gp68 from limited proteolysis by DNA templates. Both short and long templates are based on the KZP119L promoter sequence; the short template only extends to (+3) nt from the start, whereas the long template indicates a full 212 bp PCR-fragment. The full gp68 and the N-terminal fragment of gp68 extending to 382 residues are marked by arrows.

**Figure 3:**
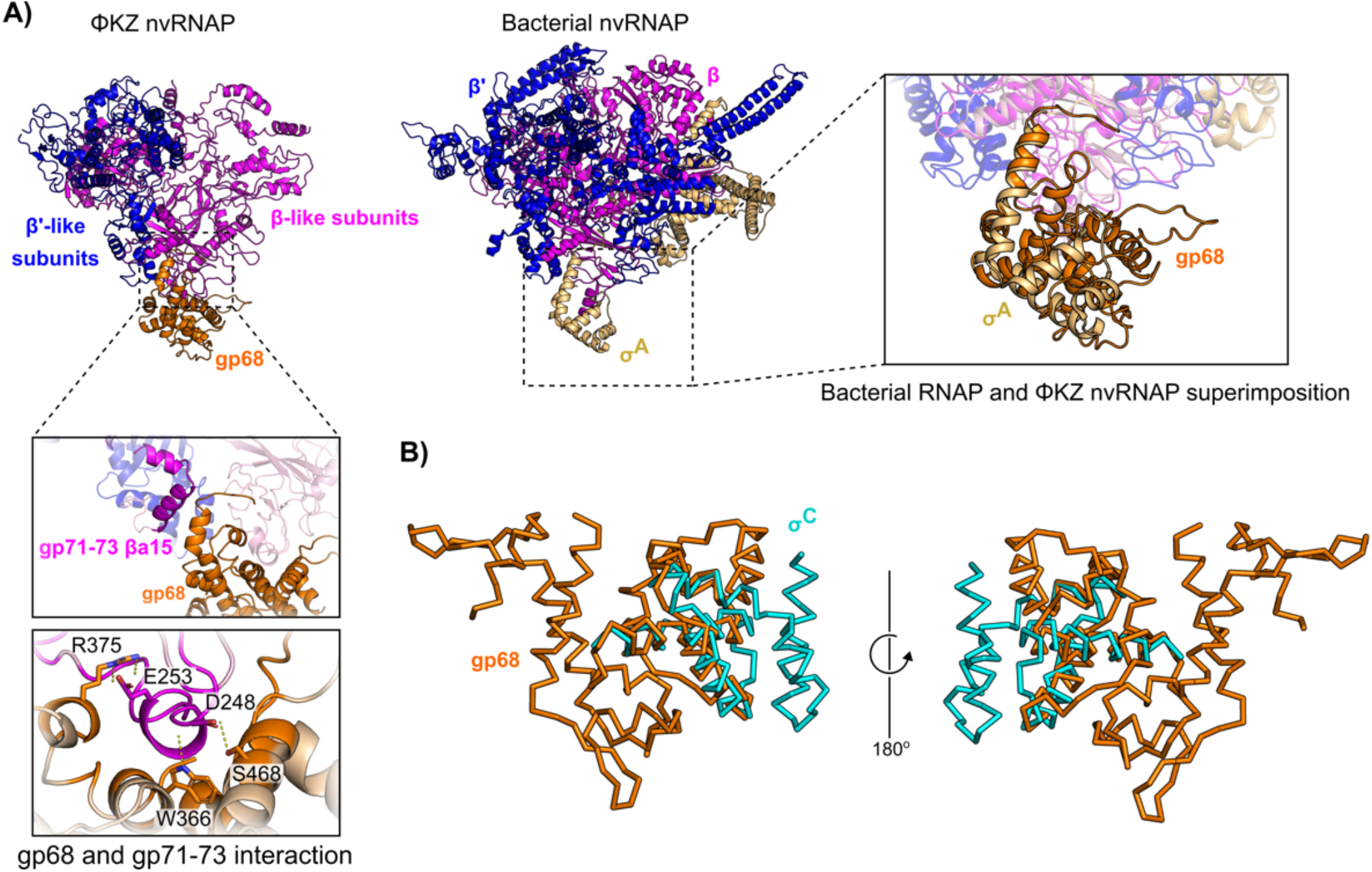
Gp68 is positioned identically to, and exhibits structural homology to, eubacterial σ-factor. a) Comparisons of *M. tuberculosis* RNAP (PDB 6EDT) and ΦKZ nvRNAP, showing the interaction surfaces between the two, and colocation of gp68 and bacterial σ^A^ factor. b) Superimposition of *M. tuberculosis* σ^C^ (PDB 2O7G) on nvRNAP gp68 showing weak structural homology identified using DALI.

Superimposition of the nvRNAP structure with that of *M. tuberculosis* RNAP revealed that the C-terminus of gp68 is located in the same position within the complex as the C-terminus of σ factor (Figure 3A). Furthermore, DALI analysis of the gp68 three-dimensional structure showed that it has a weakly similar fold to eubacterial σ factors, with the highest homology to the crystal structure of *M. tuberculosis* σ^C^ (PDB 207G)(21). Superimposition of gp68 on *M. tuberculosis* σ^C^ shows overlap of three α-helices with a RMSD of 3 Å (Figure 3B). The gp68 C-terminus is substantially expanded in comparison to the C-terminus of eubacterial σ factor, making a substantially larger interaction (2 100 vs. 1 495 Å^2^), and anchoring gp68 permanently within the complex. Previously, we have shown that the absence of gp68 compromises the assembly of a full complex, as well as the binding of the enzyme to DNA-templates and transcription initiation (Figure 2B)(19). To determine which subunits are involved in DNA binding, we carried out DNA binding assays with ΦKZ nvRNAP. We established binding by electrophoretic mobility shift assay, and then used UV-crosslinking to covalently link the bound DNA to the adjacent proteins. We subsequently SDS treated the samples to separate the polypeptides and crosslinked DNA within the complex, and separated the resulting mixture by electrophoresis, allowing the identification of several DNA-linked breakdown intermediates. Mass spectrometry identified gp123 and gp68 within the band corresponding to the fully denatured complex containing the DNA binding subunits, confirming the involvement of gp68 in DNA binding (Figure 2C). Based upon this observed involvement of gp68 in DNA binding, we then carried out limited proteolysis with two different DNA templates, a short junction DNA containing promoter, and a long dsDNA. In the absence of DNA, limited proteolysis released the N-terminus of gp68 (1-382). The presence of the short junction DNA was not protective, however the presence of the longer dsDNA was sufficient to protect gp68 from proteolysis (Figure 2D), confirming the crucial role of the fifth subunit in DNA binding, and congruent with σ factor-like activity, in that DNA long enough to allow recruitment to the polymerase active site resulted in a conformational change to a more compact complex.

Overall, our results imply that ΦKZ nvRNAP has evolved a divergent, minimal, and elegant structure to allow the assembly of a functional RNAP without the canonical α and ω subunits, and with an embedded subunit involved intimately in both the assembly of the catalytically active enzyme and DNA recruitment, that most likely began as an ancestral σ-factor, the exchange of which for alternate promoter selection became unnecessary for ΦKZ’s lifecycle. There are only a few exceptional bacteriophages known to encode multi-subunit RNAPs, including bacteriophages PBS2 (9, 26) and AR9 (27). AR9 also encodes distant homologues to β and β’ subunits whose gene products assemble into two msRNAPs (27), and a fifth subunit, gp226, crucial for promoter recognition (11), suggesting a similar mechanism to ΦKZ nvRNAP, however the AR9 complex remains catalytically active in the absence of the fifth subunit, which suggests a less intimate integration into the msRNAP complex than has been achieved by ΦKZ nvRNAP.

The conservation of the key elements of the eubacterial catalytic core in ΦKZ implies that processive polymerisation and catalysis will proceed similarly; ΦKZ has evolved an extremely streamlined msRNAP, which has no reliance on α or ω subunits, or indeed any additional stabilising contacts in their place within the complex, however the nuts and bolts of the processive polymerase are all retained as in their eubacterial orthologues. The assembly and stabilisation of the active site are likely to be divergent, however, as our biochemical results imply both to be gp68 dependent. Given the extreme divergence of gp68 from σ factor and its embedding within the complex, promoter recognition and bubble formation are also unlikely to proceed identically. Further studies of the ΦKZ nvRNAP will be required to understand how these key roles are achieved.

## Data Availability

The cryo-EM density map resolved for the ΦKZ nvRNAP complex has been deposited in the EM Databank under accession code EMD-12370, while the corresponding molecular model has been deposited in the protein data bank as PDB-ID 7NJ2.

## Acknowledgements

The authors would like to thank the Imperial Centre for Structural Biology for access to electron microscopy equipment and in particular to Paul Simpson for technical support. Data were collected at the London Consortium Electron Microscopy Facility at The Francis Crick Institute funded by Wellcome Trust Grant 206175/Z/17/Z, and would like to thank Nora Cronin for technical support. The authors are also thankful to the Center of Shared Usage «The analytical centre of nano- and biotechnologies of SPbPU» for access to scientific equipment.

## Author contributions

CHSA and MY conceived of the project. AL, MO, and MY cloned, expressed and purified proteins and carried out biochemical experiments. TA carried out mass-spectrometry experiments. ASM and PS carried out cross-linking experiments. CHSA, KR, and NdMG, prepared samples and collected electron microscopy data. CHSA, NdMG, and YTELWL, reconstructed and modelled molecular structures. All authors were involved in the interpretation of experimental results and the drafting of the manuscript.

## REFERENCES

1. Krylov, V.N., Dela Cruz, D.M., Hertveldt, K. and Ackermann, H.W. (2007) ‘φKZ-like viruses’, a proposed new genus of myovirus bacteriophages. Arch. Virol., 152, 1955–1959.

2. Krylov, V.N. and Zhazykov Zh., I. (1978) Pseudomonas bacteriophage phiKZ as a model for studying genetical control of morphogenesis. Genetika, 14, 678–685.

3. Pires, D.P., Vilas Boas, D., Sillankorva, S. and Azeredo, J. (2015) Phage Therapy: a Step Forward in the Treatment of Pseudomonas aeruginosa Infections. J. Virol., 89, 7449–7456.

4. Med Sci, T.J., Can, K., Aksu, U. and Şadi Yenen, O. (2018) Investigation of PhiKZ phage therapy against Pseudomonas aeruginosa in mouse pneumonia model. Turkish J. Med. Sci., 48, 670–678.

5. Mesyanzhinov, V. V., Robben, J., Grymonprez, B., Kostyuchenko, V.A., Bourkaltseva, M. V., Sykilinda, N.N., Krylov, V.N. and Volckaert, G. (2002) The genome of bacteriophage φKZ of Pseudomonas aeruginosa. J. Mol. Biol., 317, 1–19.

6. Mendoza, S.D., Nieweglowska, E.S., Govindarajan, S., Leon, L.M., Berry, J.D., Tiwari, A., Chaikeeratisak, V., Pogliano, J., Agard, D.A. and Bondy-Denomy, J. (2020) A bacteriophage nucleus-like compartment shields DNA from CRISPR nucleases. Nature, 577, 244–248.

7. Danilova, Y.A., Belousova, V. V., Moiseenko, A. V., Vishnyakov, I.E., Yakunina, M. V. and Sokolova, O.S. (2020) Maturation of Pseudo-Nucleus Compartment in P. aeruginosa, Infected with Giant phiKZ Phage. Viruses, 12, 1197.

8. Ceyssens, P.-J., Minakhin, L., Van den Bossche, A., Yakunina, M., Klimuk, E., Blasdel, B., De Smet, J., Noben, J.-P., Blasi, U., Severinov, K., et al. (2014) Development of Giant Bacteriophage KZ Is Independent of the Host Transcription Apparatus. J. Virol., 88, 10501–10510.

9. Clark, S., Losick, R. and Pero, J. (1974) New RNA polymerase from Bacillus subtilis infected with phage PBS2. Nature, 252, 21–24.

10. Yakunina, M., Artamonova, T., Borukhov, S., Makarova, K.S., Severinov, K. and Minakhin, L. (2015) A non-canonical multisubunit RNA polymerase encoded by a giant bacteriophage. Nucleic Acids Res., 43.

11. Sokolova, M., Borukhov, S., Lavysh, D., Artamonova, T., Khodorkovskii, M. and Severinov, K. (2017) A non-canonical multisubunit RNA polymerase encoded by the AR9 phage recognizes the template strand of its uracil-containing promoters. Nucleic Acids Res., 45, 5958–5967.

12. Iyer, L.M., Koonin, E. V. and Aravind, L. (2003) Evolutionary connection between the catalytic subunits of DNA-dependent RNA polymerases and eukaryotic RNA-dependent RNA polymerases and the origin of RNA polymerases. BMC Struct. Biol., 3, 1–23.

13. Zaychikov, E., Martin, E., Denissova, L., Kozlov, M., Markovtsov, V., Kashlev, M., Heumann, H., Nikiforov, V., Goldfarb, A. and Mustaev, A. (1996) Mapping of catalytic residues in the RNA polymerase active center. Science (80-.)., 273, 107–108.

14. Zhang, G., Campbell, E.A., Minakhin, L., Richter, C., Severinov, K. and Darst, S.A. (1999) Crystal structure of thermus aquaticus core RNA polymerase at 3.3 å resolution. Cell, 98, 811–824.

15. Heyduk, T., Heyduk, E., Severinov, K., Tang, H. and Ebright, R.H. (1996) Determinants of RNA polymerase α subunit for interaction with β, β′, and σ subunits: Hydroxyl-radical protein footprinting. Proc. Natl. Acad. Sci. U. S. A., 93, 10162–10166.

16. Minakhin, L., Bhagat, S., Brunning, A., Campbell, E.A., Darst, S.A., Ebright, R.H. and Severinov, K. (2001) Bacterial RNA polymerase subunit ω and eukaryotic polymerase subunit RPB6 are sequence, structural, and functional homologs and promote RNA polymerase assembly. Proc. Natl. Acad. Sci. U. S. A., 98, 892–897.

17. Lonetto, M., Gribskov, M. and Gross, C.A. (1992) The σ70 family: Sequence conservation and evolutionary relationships. J. Bacteriol., 174, 3843–3849.

18. Lane, W.J. and Darst, S.A. (2010) Molecular Evolution of Multisubunit RNA Polymerases: Sequence Analysis. J. Mol. Biol., 395, 671–685.

19. Orekhova, M., Koreshova, A., Artamonova, T., Khodorkovskii, M. and Yakunina, M. (2019) The study of the phiKZ phage non-canonical non-virion RNA polymerase. Biochem. Biophys. Res. Commun., 511, 759–764.

20. Glyde, R., Ye, F., Jovanovic, M., Kotta-Loizou, I., Buck, M. and Zhang, X. (2018) Structures of Bacterial RNA Polymerase Complexes Reveal the Mechanism of DNA Loading and Transcription Initiation. Mol. Cell, 70, 1111-1120.e3.

21. Thakur, K.G., Joshi, A.M. and Gopal, B. (2007) Structural and biophysical studies on two promoter recognition domains of the extra-cytoplasmic function σ factor σC from Mycobacterium tuberculosis. J. Biol. Chem., 282, 4711–4718.

22. Yokoyama, S., Yokoyama, S., Vassylyeva, M.N. and Yokoyama, S. (2002) Crystal structure of a bacterial RNA polymerase holoenzyme at 2.6. Å resolution. Nature, 417, 712–719.

23. Wang, D., Bushnell, D.A., Westover, K.D., Kaplan, C.D. and Kornberg, R.D. (2006) Structural Basis of Transcription: Role of the Trigger Loop in Substrate Specificity and Catalysis. Cell, 127, 941–954.

24. Boyaci, H., Chen, J., Jansen, R., Darst, S.A. and Campbell, E.A. (2019) Structures of an RNA polymerase promoter melting intermediate elucidate DNA unwinding. Nature, 565, 382–385.

25. Lane, W.J. and Darst, S.A. (2010) Molecular Evolution of Multisubunit RNA Polymerases: Structural Analysis. J. Mol. Biol., 395, 686–704.

26. Clark, S. (1978) Transcriptional specificity of a multisubunit RNA polymerase induced by Bacillus subtilis bacteriophage PBS2. J. Virol., 25.

27. Lavysh, D., Sokolova, M., Minakhin, L., Yakunina, M., Artamonova, T., Kozyavkin, S., Makarova, K.S., Koonin, E. V. and Severinov, K. (2016) The genome of AR9, a giant transducing Bacillus phage encoding two multisubunit RNA polymerases. Virology, 495, 185–196.

28. Tang, G., Peng, L., Baldwin, P.R., Mann, D.S., Jiang, W., Rees, I. and Ludtke, S.J. (2007) EMAN2: An extensible image processing suite for electron microscopy. J. Struct. Biol., 157, 38–46.

29. Rohou, A. and Grigorieff, N. (2015) CTFFIND4: Fast and accurate defocus estimation from electron micrographs. J. Struct. Biol., 192, 216–221.

30. Scheres, S.H.W. (2012) RELION: Implementation of a Bayesian approach to cryo-EM structure determination. J. Struct. Biol., 180, 519–530.

31. Ludtke, S.J., Baldwin, P.R. and Chiu, W. (1999) EMAN: Semiautomated Software for High-Resolution Single-Particle Reconstructions. J. Struct. Biol., 128, 82–97.

32. Drozdetskiy, A., Cole, C., Procter, J. and Barton, G.J. (2015) JPred4: a protein secondary structure prediction server. Nucleic Acids Res., 43, W389–W394.

